# A transferable IncC-IncX3 hybrid plasmid co-carrying *bla*_NDM-4_, *tet*(X4), and *tmexCD3-toprJ3* confers resistance to carbapenem and tigecycline

**DOI:** 10.1101/2021.06.30.450641

**Authors:** Aki Hirabayashi, Trung Duc Dao, Taichiro Takemura, Futoshi Hasebe, Le Thi Trang, Nguyen Ha Thanh, Hoang Huy Tran, Keigo Shibayama, Ikuro Kasuga, Masato Suzuki

## Abstract

Tigecycline is a last-resort antimicrobial that exhibits promising activity against carbapenemase-producing Enterobacterales (CPE). However, mobile tigecycline resistance genes, *tet*(X) and *tmexCD-toprJ*, have emerged in China and have spread possibly worldwide. Tet(X) family proteins, Tet(X3) to Tet(X14), function as tigecycline-inactivating enzymes, and TMexCD-TOprJ complexes function as efflux pumps for tigecycline. Here, we report a CPE isolate co-harboring both emerging tigecycline resistance factors for the first time. A carbapenem- and tigecycline-resistant *Klebsiella aerogenes* NUITM-VK5 was isolated from an urban drainage in Vietnam in 2021 and a plasmid pNUITM-VK5_mdr co-carrying *tet*(X4) and *tmexCD3-toprJ3* along with the carbapenemase gene *bla*_NDM-4_ was identified in NUITM-VK5. pNUITM-VK5_mdr was transferred to *Escherichia coli* by conjugation and simultaneously conferred high-level resistance against multiple antimicrobials, including carbapenems and tigecycline. An efflux pump inhibitor canceled TMexCD3-TOprJ3-mediated tigecycline resistance, suggesting that both tigecycline resistance factors independently and additively contribute to the high-level resistance. The plasmid had the IncX3 and IncC replicons and was estimated to be a hybrid of plasmids with different origins. Unlike IncX3 plasmids, IncC plasmids are stably maintained in an extremely broad range of bacterial hosts in humans, animals, and environment. Thus, future global spread of multidrug-resistance plasmids such as pNUITM-VK5_mdr poses a public health crisis.

## Material and methods

### Bacterial isolation and antimicrobial susceptibility testing

A carbapenem- and tigecycline-resistant *Klebsiella aerogenes* NUITM-VK5 was isolated from Kim-Nguu river in Hanoi, Vietnam in March 2021. Environmental water samples were collected and cultured using Luria-Bertani (LB) broth containing 4 mg/L of meropenem at 37°C overnight, and then further selected and isolated using CHROMagar COL-APSE (CHROMagar Microbiology) containing 4 mg/L of tigecycline. Bacterial species identification was performed using MALDI Biotyper (Bruker). Antimicrobial susceptibility testing (AST) using *Escherichia coli* ATCC 25922 as quality control was performed with agar dilution (other than colistin) and broth dilution methods (for colistin) according to the Clinical and Laboratory Standards Institute (CLSI) 2020 guidelines^12^. The categorization as susceptible (S), intermediate (I), and resistant (R) was determined according to the minimum inhibitory concentration (MIC) breakpoints. For tigecycline, AST was additionally performed in the presence or absence of 75 mg/L of the efflux pump inhibitor 1-(1-naphthylmethyl)-piperazine (NMP).

### Whole-genome sequencing and subsequent bioinformatics analysis

Whole-genome sequencing of *K. aerogenes* NUITM-VK5 was performed using MiSeq (Illumina) with MiSeq Reagent Kit v2 (300-cycle) and MinION (Oxford Nanopore Technologies) with the R9.4.1 flow cell. The library for Illumina sequencing (paired-end, insert size of 300-800 bp) was prepared using Nextera XT DNA Library Prep Kit and the library for MinION sequencing was prepared using Rapid Barcoding Kit (SQK-RBK004). Illumina reads were assembled de novo using Shovill v1.1.0 (https://github.com/tseemann/shovill) with default parameters, resulting in the draft genome (accession no.: <u>BPFV01000000</u>). MinION reads were basecalled using Guppy v4.2.2 with the high-accuracy mode and were assembled de novo using Canu v2.1.1 (https://github.com/marbl/canu) with default parameters. The overlap region in the assembled contig was detected using LAST (http://last.cbrc.jp) and was trimmed manually. Sequencing errors were corrected by Racon v1.4.13 (https://github.com/isovic/racon) twice with default parameters using MinION reads, and then corrected by Pilon v1.20.1 (https://github.com/broadinstitute/pilon/wiki) twice with default parameters using Illumina reads, resulting in a circular plasmid pNUITM-VK5_mdr (accession no.: <u>LC633285</u>).

Genome and plasmid sequences were annotated using the DFAST server (https://dfast.nig.ac.jp). Sequence type (ST) by multilocus sequence typing (MLST) analysis was determined according to the PubMLST protocol and database (https://pubmlst.org/organisms/klebsiella-aerogenes). Plasmid replicon type and antimicrobial resistance (AMR) genes were detected using PlasmidFinder v2.1 and ResFinder v4.1 with default parameters, respectively, on the CGE server (http://www.genomicepidemiology.org). Type IV secretion system (T4SS)-associated genes involved in conjugation were detected by TXSScan v1.0 (https://github.com/macsy-models/TXSS) with default parameters. Mobile gene elements (MGEs) were detected using BLAST with the ISfinder database updated on Oct 2020 (https://github.com/thanhleviet/ISfinder-sequences). Circular representation of plasmid was visualized using the CGView server (http://cgview.ca). Linear comparison of sequence alignment was performed using BLAST and visualized by Easyfig v.2.2.2 (http://mjsull.github.io/Easyfig/).

### Bacterial conjugation assay

A bacterial conjugation assay was performed as follows. LB broth cultures of the donor *K. aerogenes* NUITM-VK5 and the recipient azide-resistant *E. coli* J53 (*F*^−^*met pro Azi*^r^) were mixed in a 1:10 ratio, spotted onto MacConkey agar, and then incubated at 37°C overnight. Subsequently, the mixed cells, including transconjugants, were suspended in LB broth and then plated onto MacConkey agar containing 1 mg/L of tigecycline and 100 mg/L of sodium azide after 10-fold serial dilution, and incubated at 37°C overnight. AMR genes, *bla*_NDM_, *tet*(X), and *tmexCD-toprJ*, of transconjugants were detected by colony PCR using the following primer sets. NDM_F: GGTTTGGCGATCTGGTTTTC, NDM_R: CGGAATGGCTCATCACGATC, tetX_F: CCCGAAAATCGWTTTGACAATCCTG, tetX_R: GTTTCTTCAACTTSCGTGTCGGTAAC, tmexC_F: TGGCGGGGATCGTGCTCAAGCGCAC, tmexC_R: CAGCGTGCCCTTGCKCTCGATATCG.

## Introduction

Tigecycline, a semisynthetic glycylcycline, is considered a last-resort antimicrobial against infections caused by multidrug-resistant (MDR) gram-negative bacteria, including carbapenemase-producing Enterobacterales (CPE)^1, 2^. Carbapenemases genes, including *bla*_NDM_, *bla*_KPC_, *bla*_IMP_, *bla*_VIM_, and *bla*_OXA-48_, are often carried on plasmids, which are self-transmissible via bacterial conjugation^3^. Recently, mobile tigecycline resistance genes, *tet*(X3), *tet*(X4), and other variants, *tet*(X5) to *tet*(X14), encoding flavin-dependent monooxygenases that catalyze tigecycline degradation were emerged in Enterobacterales and *Acinetobacter* species in China and other countries^4–6^. Furthermore, a mobile tigecycline resistance gene cluster, *tmexCD-toprJ*, encoding the resistance– nodulation–cell division (RND) efflux pump that excretes multiple antimicrobials, such as tetracyclines including tigecycline, cephalosporins, fluoroquinolones, and aminoglycosides, emerged predominantly in Enterobacterales: *tmexCD1-toprJ1* was identified in plasmids in *Klebsiella* species isolated from humans and livestock in China and Vietnam^7–9^. *tmexCD2-toprJ2* was identified in the plasmid and chromosome of *Raoultella ornithinolytica* isolated from a human in China^10^. *tmexCD3-toprJ3* was identified in the chromosome of *Proteus mirabilis* isolated from livestock feces in China^11^. In this study, we identified a CPE isolate co-harboring both mobile tigecycline resistance genes, *tet*(X) and *tmexCD-toprJ*, for the first time and characterized a transferable IncC-IncX3 hybrid plasmid co-carrying *bla*_NDM-4_, *tet*(X4), and *tmexCD3-toprJ3* in *K. aerogenes* isolated in Vietnam.

## Results and discussion

A carbapenem- and tigecycline-resistant *K. aerogenes* isolate NUITM-VK5 was obtained from an urban drainage in Hanoi, Vietnam, in March 2021. *K. aerogenes* (former *Enterobacter aerogenes*) is an important human opportunistic pathogen and a frequent cause of nosocomial infections. The result of AST of *K. aerogenes* NUITM-VK5 showed that NUITM-VK5 was resistant to almost all antimicrobials tested (Table 1). The MICs of tigecycline, tetracyclines, carbapenems, cephalosporins, fluoroquinolone, and aminoglycosides (other than amikacin) were more than 128 mg/L (R) and that of colistin was 4 mg/L (R), whereas that of amikacin was 32 mg/L (I).

**Table 1.**
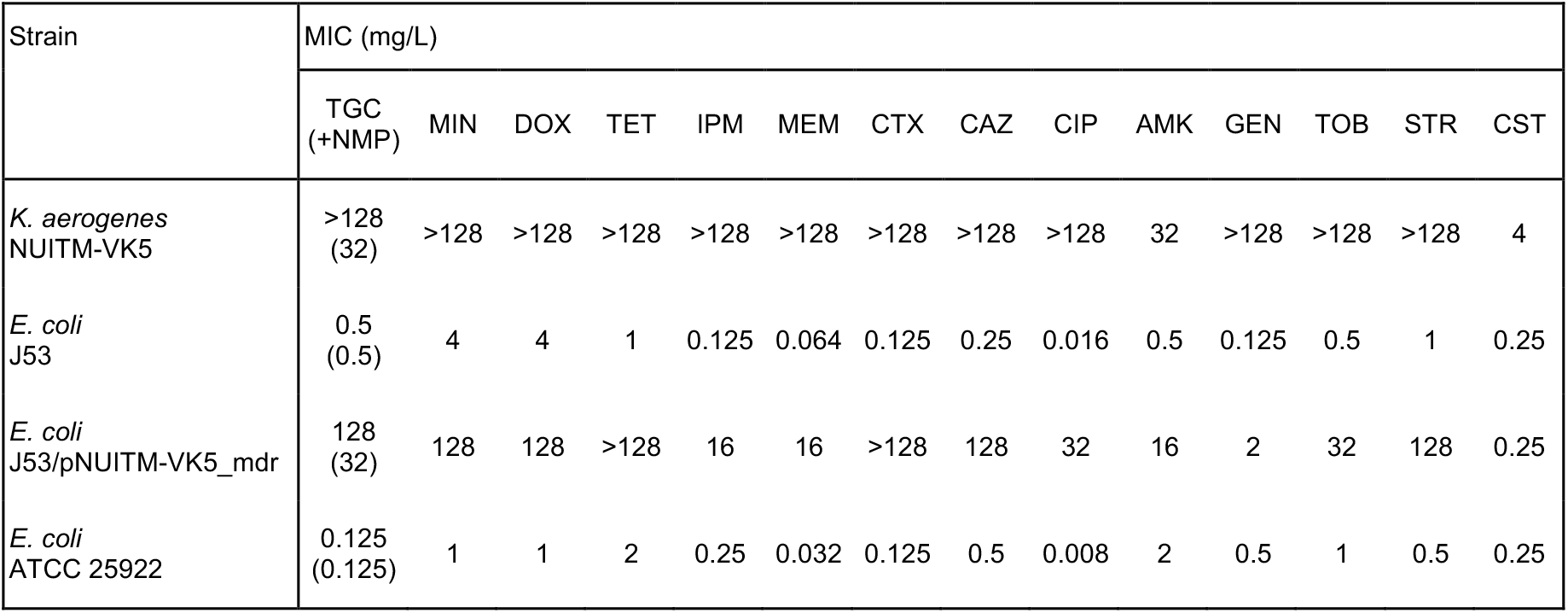
Minimum inhibitory concentrations (MICs) of antimicrobials against *K. aerogenes* NUITM-VK5 and its transconjugant of *E. coli* J53 harboring the plasmid pNUITM-VK5_mdr (J53/pNUITM-VK5_mdr). The efflux pump inhibitor 1-(1-naphthylmethyl)-piperazine (NMP) was used at 75 mg/L. TGC, tigecycline; MIN, minocycline; DOX, doxycycline; TET, tetracycline; IPM, imipenem; MEM, meropenem; CTX, cefotaxime; CAZ, ceftazidime; CIP, ciprofloxacin; AMK, amikacin; GEN, gentamicin; TOB, tobramycin; STR, streptomycin; CST, colistin.

| Strain | MIC (mg/L) |  |  |  |  |  |  |  |  |  |  |  |  |  |
| --- | --- | --- | --- | --- | --- | --- | --- | --- | --- | --- | --- | --- | --- | --- |
|  | TGC<br>(+NMP) | MIN | DOX | TET | IPM | MEM | CTX | CAZ | CIP | AMK | GEN | TOB | STR | CST |
| *K. aerogenes*<br>NUITM-VK5 | >128<br>(32) | >128 | >128 | >128 | >128 | >128 | >128 | >128 | >128 | 32 | >128 | >128 | >128 | 4 |
| *E. coli*<br>J53 | 0.5<br>(0.5) | 4 | 4 | 1 | 0.125 | 0.064 | 0.125 | 0.25 | 0.016 | 0.5 | 0.125 | 0.5 | 1 | 0.25 |
| *E. coli*<br>J53/pNUITM-VK5_mdr | 128<br>(32) | 128 | 128 | >128 | 16 | 16 | >128 | 128 | 32 | 16 | 2 | 32 | 128 | 0.25 |
| *E. coli*<br>ATCC 25922 | 0.125<br>(0.125) | 1 | 1 | 2 | 0.25 | 0.032 | 0.125 | 0.5 | 0.008 | 2 | 0.5 | 1 | 0.5 | 0.25 |

Short-read sequence analysis of *K. aerogenes* NUITM-VK5 with MiSeq constructed the draft genome consisting of 181 contigs (5.9 Mbp, accession no.: <u>BPFV01000000</u>). MLST analysis showed that NUITM-VK5 belonged to sequence type 4 (ST4). Detection of AMR genes using ResFinder v4.0 with the modified library including nucleotide sequences of known variants of *tmexCD-toprJ* revealed that NUITM-VK5 harbored *bla*_NDM-4_, *tet*(X4), and *tmexCD3-toprJ3* along with multiple clinically relevant AMR genes, such as *bla*_CTX-M-14_ (extended-spectrum β-lactamase gene), *qnrS1* (fluoroquinolone resistance gene), *aac(6′)-lb-cr* (aminoglycoside resistance gene), and *cfr* (phenicol/lincosamide resistance gene). NUITM-VK5 was colistin-resistant, but did not harbor known mobile colistin resistance genes, such as *mcr*. The coding sequences of *tet*(X4) and *tmexCD3-toprJ3* in NUITM-VK5 were highly identical to those of *tet*(X4) in *E. coli* 47EC (accession no.: <u>MK134376</u>) isolated from a pig in China in 2018 and *tmexCD3-toprJ3* in *P. mirabilis* RGF134-1 (accession no.: <u>CP066833</u>) isolated from a pig in China in 2019, respectively (Fig. S1). The identity for *tet*(X4) was 97.7% (1131/1158 nt), resulting in 12 amino acid substitutions (I356A, K359R, E363A, T366I, Q367I, I370T, K374S, P375L, T378S, Q381K, L383M, and V385L). For *tmexC3*, the identity was 99.7% (1161/1164 nt), resulting in three amino acid substitutions (Q187H, T256M, and A386T); For *tmexD3*, the identity was 99.9% (3133/3135 nt), resulting in two amino acid substitutions (V610L and L611F). For *toprJ3*, the identity was 100% (1434/1434 nt).

The subsequent long-read sequence analysis of *K. aerogenes* NUITM-VK5 with MinION successfully constructed a circular plasmid pNUITM-VK5_mdr (240.5 kbp, accession no.: <u>LC633285</u>) co-carrying the aforementioned AMR genes detected in the draft genome (Fig. 1). Detection of plasmid replicons with PlasmidFinder v2.1 revealed that pNUITM-VK5_mdr had two replicons classified to incompatibility groups C (IncC) and X3 (IncX3). *bla*_NDM-4_, *tet*(X4), and *tmexCD3-toprJ3* were encoded on different locations on pNUITM-VK5_mdr, and the GC contents of the regions surrounding those AMR genes (61.6% for *bla*_NDM-4_, 37.4% for *tet*(X4), and 66.0% for *tmexCD3-toprJ3*) were different from the average of the whole plasmid (51.4%), suggesting that these regions were acquired via horizontal gene transfer (HGT) mediated by mobile gene elements (MGEs) (Fig. 1)^7, 13^. *tmexCD3-toprJ3* was flanked by two MGEs, IS*L3* and IS*1182*, in pNUITM-VK5_mdr; this genetic structure was different from the *tmexCD3-toprJ3*-surrounding region in the chromosomal SXT/R391 integrative conjugative element (ICE) in *P. mirabilis* RGF134-1 (Fig. 1, lower)^11^. *bla*_NDM-4_ and *tet*(X4) in pNUITM-VK5_mdr were estimated to be acquired via HGT mediated by MGEs, IS*26* and IS*Vsa3*, respectively, as previously reported^14, 15^.

**Fig. 1.**
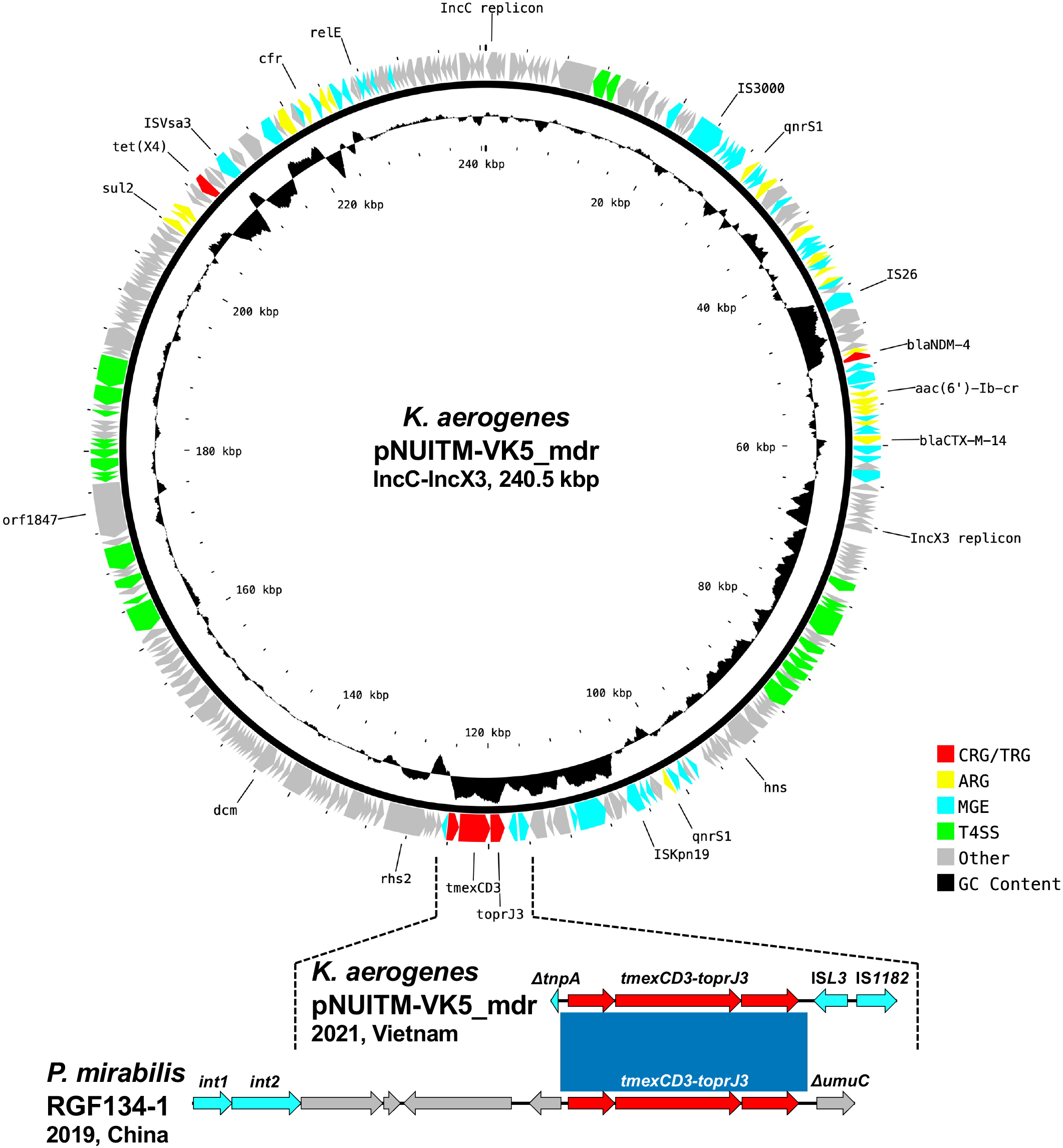
Upper: Circular representation of a 240.5-kbp IncC-IncX3 hybrid plasmid, pNUITM-VK5_mdr, co-carrying multiple antimicrobial resistance genes including *bla*_NDM-4_*, tet*(X4), and *tmexCD3-toprJ3* in *K. aerogenes* NUITM-VK5 isolated in Vietnam in 2021. Lower: Linear comparison of *tmexCD3-toprJ3*-containing regions in *K. aerogenes* pNUITM-VK5_mdr and in a chromosome of *P. mirabilis* RFG134-1 isolated in China in 2019. Red, yellow, cyan, green, gray, and black indicate carbapenem and tetracycline resistance genes (CRG/TRG), other AMR genes (ARG), mobile gene elements (MGE), type IV secretion system (T4SS)-associated genes involved in conjugation, other coding sequences (Other), and GC content, respectively. The blue color in comparison of sequences indicates almost 100% identity.

A bacterial conjugation assay using *E. coli* J53 as the recipient strain showed that *K. aerogenes* NUITM-VK5 transferred pNUITM-VK5_mdr to J53 at a frequency of 1.0 × 10^−6^ after overnight co-culture at 37°C. The transconjugant strain (J53/pNUITM-VK5_mdr) was confirmed to co-harbor *bla*_NDM-4_, *tet*(X4), and *tmexCD3-toprJ3* by PCR and was resistant to almost all antimicrobials, including carbapenems and tigecycline (Table 1). The transconjugant strain was only susceptible to colistin, although parental NUITM-VK5 was resistant, suggesting that colistin resistance of NUITM-VK5 was due to other factors, including chromosomal gene mutations, other than pNUITM-VK5_mdr. The addition of the efflux pump inhibitor NMP reduced the MIC of tigecycline from 128 mg/L or higher to 32 mg/L in NUITM-VK5 and the transconjugant strain () (Table 1). Since the MIC of tigecycline against *E. coli* J53 was 0.5 mg/L, 32 mg/L for the MIC against J53/pNUITM-VK5_mdr in the presence of NMP was still high, suggesting that TMexCD3-TOprJ3 and Tet(X4) contributed to tigecycline resistance independently and additively and Tet(X4) remained active even when the RND efflux pump was inhibited. On the other hand, the addition of NMP did not affect the MIC of meropenem against NUITM-VK5 and J53/pNUITM-VK5_mdr, indicating that TMexCD3-TOprJ3 does not contribute to carbapenem resistance (data not shown).

BLASTn analysis using megablast showed that no plasmid showed more than 90% identity in more than 80% regions with the IncC-IncX3 hybrid plasmid pNUITM-VK5_mdr in the NCBI database of Nucleotide collection (nr/nt). A comparison with the known IncC plasmids showed that the IncX3 backbone of pNUITM-VK5_mdr might include the 83.5-kb region between IS*3000* and IS*Kpn19* and the IncC backbone might include the remaining region (Fig. 1). In this case, *bla*_NDM-4_ and *tet*(X4) were derived from the IncX3 and IncC backbones, respectively, and *tmexCD3-toprJ3* located at the boundary of both backbones. IncC (former IncA/C2) is divided into type 1 and type 2^16, 17^. The IncC backbone of pNUITM-VK5_mdr would belong to type 2 as it had *rhs2* and *orf1847*, which are characteristic genetic makers of type 2 (Fig. 1).

IncC, which is involved in the spread of AMR genes, has an extremely broad host range of Gammaproteobacteria^18^, whereas IncX3, which is also involved in the spread of AMR genes, such as *bla*_NDM_, has a narrow host range of Enterobacterales^19^. The combination of two incompatibility groups resulted in an IncC-IncX3 hybrid plasmid, which is expected to possess an increased risk of carrying more AMR genes and spreading more stably and efficiently among various bacterial species in humans, animals, and environment. Moreover, the acquisition of an additional RND efflux pump TMexCD3-TOprJ3 could allow the bacterial host to survive in a variety of conditions such as antimicrobial exposure, leading to further accumulation of AMR genes into the host genome via HGT^20^.

## Conclusions

The future global spread of such a broad-host-range self-transferable MDR plasmid among human pathogens poses a public health crisis and needs to be continuously monitored according to the One-Health approach.

## Transparency declarations

None to declare.

---

**Fig. S1.**

Multiple sequence alignment analyzed by MAFFT v7.480. (A) Comparison between gene products of *tet*(X4) in *K. aerogenes* NUITM-VK5 (accession no.: <u>BPFV01000000</u>) and the reference gene in *E. coli* 47EC (accession no.: <u>MK134376</u>). Twelve amino acid substitutions (I356A, K359R, E363A, T366I, Q367I, I370T, K374S, P375L, T378S, Q381K, L383M, and V385L) were found. (B) Comparison between gene products of *tmexC3* in *K. aerogenes* NUITM-VK5 and the reference sequence in *P. mirabilis* RGF134-1 (accession no.: <u>CP066833</u>). Three amino acid substitutions (Q187H, T256M, and A386T) were found. (C) Comparison between gene products of *tmexD3* in *K. aerogenes* NUITM-VK5 and the reference sequence in *P. mirabilis* RGF134-1. Two amino acid substitutions (V610L and L611F) were found.

