## Supplementary material for "A transferable IncC-IncX3 hybrid plasmid co-carrying *bla*_NDM-4_, *tet*(X4), and *tmexCD3-toprJ3* confers resistance to carbapenem and tigecycline": Fig. S1

## A

### Tet(X4)

```
BPFV01000000   MSNKEKQMNLLSDKNVAIIGGGPVGLTMAKLLQQNGIDVSVYERDNDREARIFGGTLDLH
MK134376       MSNKEKQMNLLSDKNVAIIGGGPVGLTMAKLLQQNGIDVSVYERDNDREARIFGGTLDLH
               ************************************************************

BPFV01000000   KGSGQEAMKKAGLLQTYYDLALPMGVNIADEKGNILSTKNVKPENRFDNPEINRNDLRAI
MK134376       KGSGQEAMKKAGLLQTYYDLALPMGVNIADEKGNILSTKNVKPENRFDNPEINRNDLRAI
               ************************************************************

BPFV01000000   LLNSLENDTVIWDRKLVMLEPGKKKWTLTFENKPSETADLVIIANGGMSKVRKFVTDTEV
MK134376       LLNSLENDTVIWDRKLVMLEPGKKKWTLTFENKPSETADLVIIANGGMSKVRKFVTDTEV
               ************************************************************

BPFV01000000   EETGTFNIQADIHHPEVNCPGFFQLCNGNRLMAAHQGNLLFANPNNNGALHFGISFKTPD
MK134376       EETGTFNIQADIHHPEVNCPGFFQLCNGNRLMAAHQGNLLFANPNNNGALHFGISFKTPD
               ************************************************************

BPFV01000000   EWKNQTQVDFQNRNSVVDFLLKEFSDWDERYKELIRVTSSFVGLATRIFPLGKSWKSKRP
MK134376       EWKNQTQVDFQNRNSVVDFLLKEFSDWDERYKELIRVTSSFVGLATRIFPLGKSWKSKRP
               ************************************************************

BPFV01000000   LPITMIGDAAHLMPPFAGQGVNSGLMDALILSDNLTNGKFNSIEEAIENYEQQMFAYGRE
MK134376       LPITMIGDAAHLMPPFAGQGVNSGLMDALILSDNLTNGKFNSIEEAIENYEQQMFIYGKE
               ******************************************************* **:*

BPFV01000000   AQAESIINETEMFSLDFSFQKLMNL
MK134376       AQEESTQNEIEMFKPDFTFQQLLNV
               ** **  ** ***. **:**:*:*:
```

## B

### TMexC3

```
BPFV01000000   MNKFREWITFSVISCLVAVTLVGCDKPEEQGEEAPAREVDVLSVQTEPFTVVAELPGRIE
CP066833       MNKFREWITFSVISCLVAVTLVGCDKPEEQGEEAPAREVDVLSVQTEPFTVVAELPGRIE
               ************************************************************

BPFV01000000   PVRVAEVRARVAGIVLKRTFEEGADVKAGDVLFQIDPAPFKAALSRAQGELARAEAQLFQ
CP066833       PVRVAEVRARVAGIVLKRTFEEGADVKAGDVLFQIDPAPFKAALSRAQGELARAEAQLFQ
               ************************************************************

BPFV01000000   AQAMVRRYEPLVKINAVSQQDFDNAKAALQSAQADKRSAQANVETARLDLGYAEVRAPIA
CP066833       AQAMVRRYEPLVKINAVSQQDFDNAKAALQSAQADKRSAQANVETARLDLGYAEVRAPIA
               ************************************************************

BPFV01000000   GRIGRAHVTEGALVGQGEATLLARIQQLDPVYADFTQPAADALRLRAAIAEGKVAGASDQ
CP066833       GRIGRAQVTEGALVGQGEATLLARIQQLDPVYADFTQPAADALRLRAAIAEGKVAGASDQ
               ******:*****************************************************

BPFV01000000   PLSLRVDGTDIESKGMLLFTDISVDRSTGQIALRGQFDNPEGVLLPGMYVRVRTPQGLNQ
CP066833       PLSLRVDGTDIESKGTLLFTDISVDRSTGQIALRGQFDNPEGVLLPGMYVRVRTPQGLNQ
               **************** *******************************************

BPFV01000000   NAILVPQRAVQRSADGQASVMLLGEGDTVEVRQVTTGAMQGSRWQISEGLQAGDKVITSS
CP066833       NAILVPQRAVQRSADGQASVMLLGEGDTVEVRQVTTGAMQGSRWQISEGLQAGDKVITSS
               ************************************************************

BPFV01000000   LAAIRPGAKVIPREQGAAEKAPQSQTQ
CP066833       LAAIRPGAKVIPREQGAAEKAPQSQAQ
               *************************:*
```

## C

### TMexD3

```
BPFV01000000   MPLFFIRRPNFAWVVALFISLGGLLVIPFLPVAQYPNVAPPQITVTATYPGASAQVLTDS
CP066833       MPLFFIRRPNFAWVVALFISLGGLLVIPFLPVAQYPNVAPPQITVTATYPGASAQVLTDS
               ************************************************************

BPFV01000000   VTSVIEEELNGAKNLLYFESTSNANGIAEITVTFQPGTDPELAQVDVQNRLKKAEARMPQ
CP066833       VTSVIEEELNGAKNLLYFESTSNANGIAEITVTFQPGTDPELAQVDVQNRLKKAEARMPQ
               ************************************************************

BPFV01000000   AVLTLGIQTEQATAGFLLIYALSYTDGDKDSDVTALADYAARSINNEIRRVPGVGKLQFF
CP066833       AVLTLGIQTEQATAGFLLIYALSYTDGDKDSDVTALADYAARSINNEIRRVPGVGKLQFF
               ************************************************************

BPFV01000000   ASEAAMRVWIDPQKLVGYGLSIDDVNNAIRAQNVQVPAGAFGSTPGSSEQELTATLAVKG
CP066833       ASEAAMRVWIDPQKLVGYGLSIDDVNNAIRAQNVQVPAGAFGSTPGSSEQELTATLAVKG
               ************************************************************

BPFV01000000   TLDNPQEFAAIVLRANQDGSRLTLGDVARIEVGSQDYNFGSRQDGKPAVAAAVQLSPGAN
CP066833       TLDNPQEFAAIVLRANQDGSRLTLGDVARIEVGSQDYNFGSRQDGKPAVAAAVQLSPGAN
               ************************************************************

BPFV01000000   AIQTAEAVKQRLTELSANFPDNVEFSVPYDTSRFVDVAIDKVIMTLIEAMVLVFLVMFLF
CP066833       AIQTAEAVKQRLTELSANFPDNVEFSVPYDTSRFVDVAIDKVIMTLIEAMVLVFLVMFLF
               ************************************************************

BPFV01000000   LQNVRYTLIPSIVVPVCLLGTLTFMYLLGFSVNMMTMFGMVLAIGILVDDAIVVVENVER
CP066833       LQNVRYTLIPSIVVPVCLLGTLTFMYLLGFSVNMMTMFGMVLAIGILVDDAIVVVENVER
               ************************************************************

BPFV01000000   IMAEEGLAPVPATIKAMGQVSGAIIGITLVLSAVFLPLAFMAGSVGVIYQQFSLSLAVSI
CP066833       IMAEEGLAPVPATIKAMGQVSGAIIGITLVLSAVFLPLAFMAGSVGVIYQQFSLSLAVSI
               ************************************************************

BPFV01000000   LFSGFLALTFTPALCATLLKPIPVGHHEKTGFFGWFNRKFTSLTSRYTKLNDKLVPRAGR
CP066833       LFSGFLALTFTPALCATLLKPIPVGHHEKTGFFGWFNRKFTSLTSRYTKLNDKLVPRAGR
               ************************************************************

BPFV01000000   VMFIYLGVVVLMGFLYMRLPESFVPVEDQGYMIVDIQLPPGATRERTSAAGGELESFLMA
CP066833       VMFIYLGVVVLMGFLYMRLPESFVPVEDQGYMIVDIQLPPGATRERTSAAGGELESFLMA
               ************************************************************

BPFV01000000   REAVQTTFLLFGFSFSGMGENAAIAFPLLKDWSERDSSQSPEAESVAVNEHFANLDDGAI
CP066833       REAVQTTFLVLGFSFSGMGENAAIAFPLLKDWSERDSSQSPEAESVAVNEHFANLDDGAI
               *********::*************************************************

BPFV01000000   MSVPPPPIEGLGNSGGFALRLQDRAGLGRDALLAARDEVLGKVNGNPKFLYAMMEGLAEA
CP066833       MSVPPPPIEGLGNSGGFALRLQDRAGLGRDALLAARDEVLGKVNGNPKFLYAMMEGLAEA
               ************************************************************

BPFV01000000   PQLRLVIDREQARTLGVSFEAISSALSTAFGSSVINDFANAGRQQRVVVQAEQAERMTPE
CP066833       PQLRLVIDREQARTLGVSFEAISSALSTAFGSSVINDFANAGRQQRVVVQAEQAERMTPE
               ************************************************************

BPFV01000000   SVLRLHVPNDSGSLVPLSAFVTTSWEEGPVQVARYNGYPSIRIAGDAAPGVSTGEAMLEL
CP066833       SVLRLHVPNDSGSLVPLSAFVTTSWEEGPVQVARYNGYPSIRIAGDAAPGVSTGEAMLEL
               ************************************************************

BPFV01000000   ERIAAELPEGIGYEWTGLSYQERVASGQATMLFALAITVVFLLLVALYESWAIPLTVMLI
CP066833       ERIAAELPEGIGYEWTGLSYQERVASGQATMLFALAITVVFLLLVALYESWAIPLTVMLI
               ************************************************************

BPFV01000000   VPVGALGAVLAVTAIGLPNDVYFKVGLITVIGLAAKNAILIVEFAKDLWEDGYSLRDAAV
CP066833       VPVGALGAVLAVTAIGLPNDVYFKVGLITVIGLAAKNAILIVEFAKDLWEDGYSLRDAAV
               ************************************************************

BPFV01000000   EAARLRFRPIIMTSMAFMLGVVPLAIATGAGAASQRALGTGVLGGMLSATMLGVIFVPIF
CP066833       EAARLRFRPIIMTSMAFMLGVVPLAIATGAGAASQRALGTGVLGGMLSATMLGVIFVPIF
               ************************************************************

BPFV01000000   FVWVLSLLRTKPQQTDNHPLHKAE
CP066833       FVWVLSLLRTKPQQTDNHPLHKAE
               ************************
```
